# A deep intronic variant activates a pseudoexon in the *MTM1* gene in a family with X-linked myotubular myopathy

**DOI:** 10.1101/2020.05.28.122333

**Authors:** Jamie Fitzgerald, Cori Feist, Paula Dietz, Stephen Moore, Donald Basel

**Author notes:** **Corresponding Author:** Jamie Fitzgerald, Bone and Joint Center, Henry Ford Hospital, Integrative Biosciences Center, 6135 Woodward Avenue, Detroit, MI 48202, USA.

## Abstract

We report a novel intronic variant in the *MTM1* gene in four males in a family with severe X-linked myotubular myopathy. The A>G variant in deep intronic space activates a cryptic 5’ donor splice site resulting in the inclusion of a 48bp pseudoexon into the mature *MTM1* mRNA. The variant is present in all affected males, absent in unaffected males and heterozygous in the mother of the affected males. The included intronic sequence contains a premature stop codon and experiments using a translational inhibitor indicate that the mutant mRNAs undergo nonsense-mediate decay. We conclude that affected males produce no, or low, levels of myotubularin-1 protein leading to a severe neonatal myopathy. The study highlights the need to consider non-coding variants in genomic screening in families with X-linked myotubular myopathy.

## Introduction

RNA splicing is a fundamental and tightly-regulated process that directs the production of multiple protein isoforms from a single gene by removing introns and combining different exons. Pseudoexons are intronic sequences, sometimes far from exons, that have the basic consensus sequences at the 5’ and 3’ splice sites but are not normally spliced into mature mRNA (Dhir et al. 2010; Buratti et al. 2007; Dhir and Buratti 2010). From a human disease perspective, many pseudoexon intronic sequences are poised on the brink of becoming exons but require an additional sequence change to activate cryptic splice sites. When these mutations occur and the cryptic splice sites are activated, pseudoexons can be ‘spliced-in’ to become ‘pathogenic pseudoexons’ (Dhir and Buratti 2010). *De novo* mutations that activate pseudoexons have been reported in approximately 60 genes (Buratti et al. 2011).

X-linked myotubular myopathy (XLMTM) (OMIM310400) is the most common of the centronuclear myopathies and the only one to have an X-linked inheritance pattern. Patients show generalized muscle weakness and respiratory insufficiency that often leads to death in the neonatal period or childhood. *MTM1* is a 14 exon gene located on the distal long arm of the X chromosome, band Xq28 (NM_000252.2). The *MTM1* gene product, myotubularin-1, functions as a 3-phosphoinositide phosphatase involved in phosphoinositol signaling and is required for skeletal muscle cell differentiation (Spiro, Shy, and Gonatas 1966; van Wijngaarden et al. 1969). More than 400 *MTM1* mutations in XLMTM have been described to date. Pathogenic variants that result in the loss of myotubularin-1 protein are frequently associated with demise as a result of respiratory insufficiency but a number of affected individuals have survived with the advances seen in cardiopulmonary support technology. M

Here we report a novel pathogenic pseudoexon variant in the *MTM1* gene in a family with a severe form of XLMTM.

## Materials and Methods

### Cell culture and cycloheximide treatment

Primary fibroblasts were established from patient skin biopsies and maintained in DMEM/10% fetal calf serum supplemented with 100U/ml pen and 100μg/ml strep (GIBCO)(Bateman et al. 1999). To stabilize proteins containing premature stop codons, cultures were treated for 6 hrs with 100mg/ml cycloheximide and RNA isolated for RT-PCR.

### RT-PCR

Total RNA was isolated from cultures using RNeasy kit (Qiagen), quantitated and 1 μg of RNA reverse transcribed into cDNA using iScript kit (Bio-Rad). Three overlapping *MTM1* cDNAs were amplified using three pairs of cDNA primers from Tosch et al(Tosch et al. 2010) using the PrimeSTAR GxL kit (Takara). Primers are F1/R1 (ATGGCTTCTGCATCAACTTC / TGGAATTCGATTTCGGGAC) for fragment 1 (678nt), F2/R2 (GTTCCGTATCGTGCCTCAG / GGAGAACGGTCAGCATCGG) for fragment 2 (698nt) and F3/R3 (AGAATGGATAAGTTTTGGAC / TTATTTCGAGCTCTAATGCG) for fragment 3 (622nt).

For quantitative PCR, cDNAs were diluted 1/20 and amplification conducted using SYBR Green (Applied Biosystems) with the F3/R3 primer set under conditions of maximal efficiency on an ABI2400 quantitative thermocycler. Data analysis was carried out using the comparative ΔCt method using the L32 ribosomal gene as a housekeeping gene.

## Results

### Clinical description

The family was identified in 2005 when a neonatal male (IV-1 in Figure 1) presented to the neonatal intensive care unit with lack of respiratory effort and severe hypotonia. He was born to a 16-year old G1P0 woman at 36 weeks gestation. Her pregnancy was complicated by severe polyhydramnios. He required mechanical ventilation and underwent several laboratory evaluations including normal plasma creatine kinase levels, normal methylation for Prader-Willi syndrome, two copies of *SMN1*, normal testing for myotonic dystrophy type 1, and normal very long chain fatty acids. Muscle biopsy at day of life 7 was concerning for a central nuclear myopathy. Blood was sent for *MTM1* gene sequencing, which did not reveal any sequence changes in the 14 coding exons or the 5’-untranslated region of exon 1. He was transitioned to comfort care and died at day of life 25. No autopsy was performed. His maternal cousin (IV-6), also male, was born in 2009 with a similar clinical presentation including severe polyhydramnios precipitating delivery at 35w5d gestation. The baby was transferred to the neonatal intensive care unit at delivery with hypotonia and absent respiratory effort requiring mechanical ventilation. Muscle biopsy at day of life 4 revealed relative smallness of type 1 and 2b fibers but no discernable features of a centronuclear myopathy. He was transitioned to comfort care and died at 11 days of age. Autopsy was performed but of limited clinical usefulness. It was at this time that the extended family history of an X-linked congenital myopathy was identified with at least three generations of males presenting with polyhydramnios in the third trimester who were “floppy” at birth and died in the neonatal period. In 2011, another male (IV-2) was born to the obligate carrier female (III-1), with the same condition. He was delivered at 35 weeks gestation by repeat Cesarean section for severe polyhydramnios. A birth he had no spontaneous respirations and required aggressive resuscitation. He was intubated but after no response to chest compressions, fluid bolus, and epinephrine resuscitation efforts were discontinued. He died four minutes later. Blood was sent for karyotype (46,XY) but no other genetic testing or autopsy was performed. In 2016, a third affected son (IV-3) was born to III-1 by repeat Cesarean section at 37 weeks for polyhydramnios. This male was intubated at birth for no respiratory effort but was able to transition to continuous positive airway pressure at two days of age. For unclear reasons, this child appeared less severely affected and was able to transition to bilevel positive airway pressure at 14 weeks of life. He underwent G-J tube placement at 11 weeks of life as well as various diagnostic studies. Testing for SMN1-related spinal muscular atrophy, Prader-Willi syndrome, myotonic dystrophy type 1 and trio whole exome sequencing in a CLIA/CAP approved laboratory was non-diagnostic. Muscle biopsy at 11 weeks of life was suggestive of centronuclear myopathy. This child was evaluated throughout his admission by pediatric neuromuscular specialists and placed on mestinon 1mg/kg/dose TID at 15 weeks of life. He was discharged from the neonatal intensive care unit at 18 weeks of life (55 weeks corrected gestational age). He died suddenly of cardiorespiratory arrest at 1 year and 11 days of life. Pedigree of the extended family is shown in Fig. 1.

**Fig. 1:**
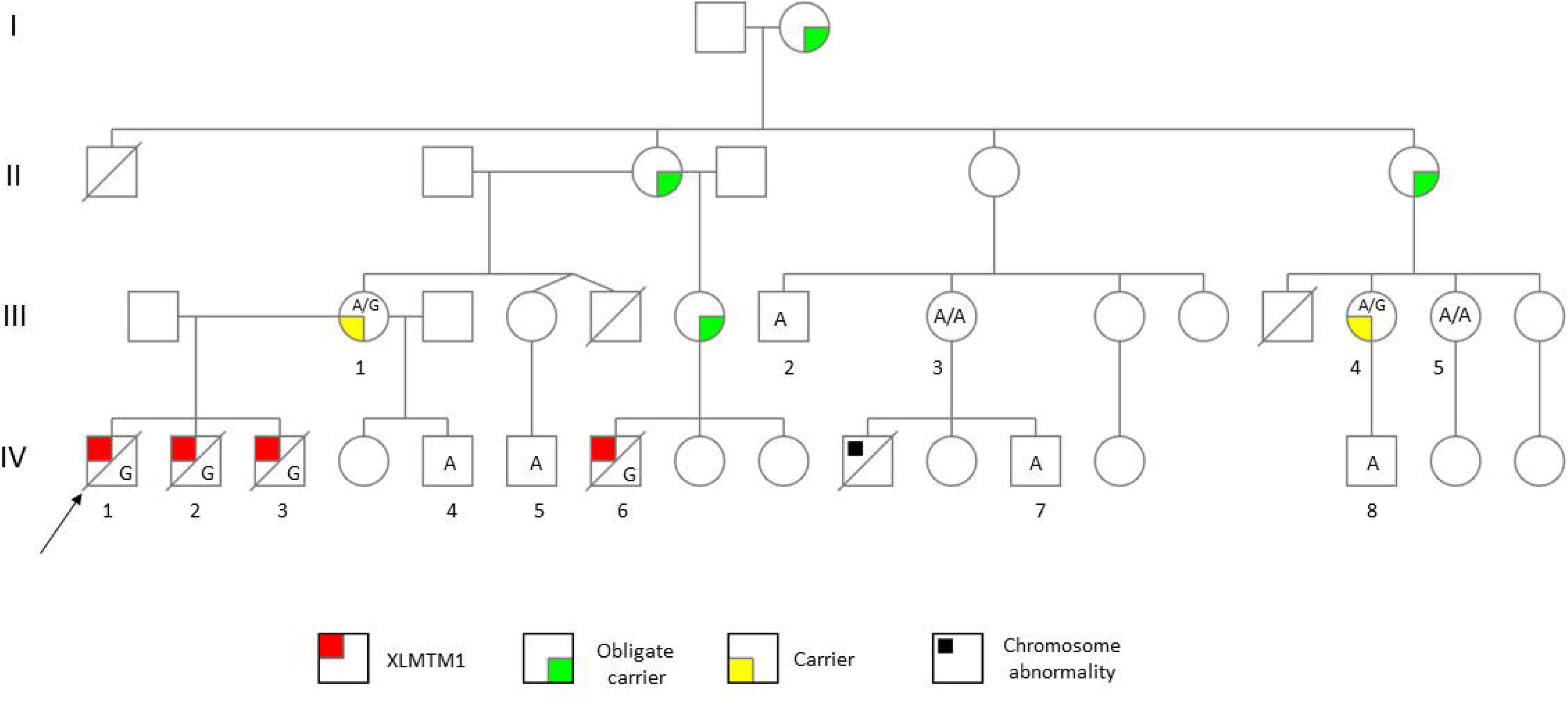
Pedigree of extended family with XL*MTM1*.

### Identification of intronic variant

Since the proband was considered to have an X-linked myopathy similar to XL*MTM1*, the coding region of the *MTM1* gene was sequenced manually independent of the clinical sequencing. Direct DNA sequencing of *MTM1* failed to identify a mutation in the *MTM1* exons and intron-exon boundaries. Since it is emerging that a significant fraction of genetic disease is due to mutations in non-coding regions, RT-PCR was conducted on RNA isolated from fibroblasts from one of the affected males (Fig. 2A). One fragment (PCR3) produced a larger than expected RT-PCR product in patient compared to control cDNA suggesting the inclusion of additional sequence in the mature mRNA. Sequencing the PCR3 product revealed that the additional sequence was a 48bp stretch of intronic sequence spanning chrX:149831277-149831324 in *MTM1* intron 13. Sequencing of genomic DNA flanking the included sequence revealed an A>G substitution 5 bases downstream of the end of the included sequence at chrX:149831329 A>G c.1468-577 (*MTM1*, NM_000252.2) (Fig. 2B). This variant is 2362 bp downstream from exon 13 and 579 bp from the start of exon 14. The variant is unique and not present in the EXAC or SNP human variant databases and no ESTs containing the included intron have been described. Genotyping thirteen family members demonstrated that all the affected (individuals IV-1, -2, -3, and -6), but none of the unaffected males (IV-4, -5, -7, and -8), carry the variant. The mother (III-1) of three affected males is heterozygous for the variant, as expected, as is the mother (III-4) of an unaffected boy (IV-8). Sequencing chromatograms of the carrier mother and affected and unaffected sons showing the pathogenic A>G variant, is shown in Fig. 2C.

**Fig. 2:**
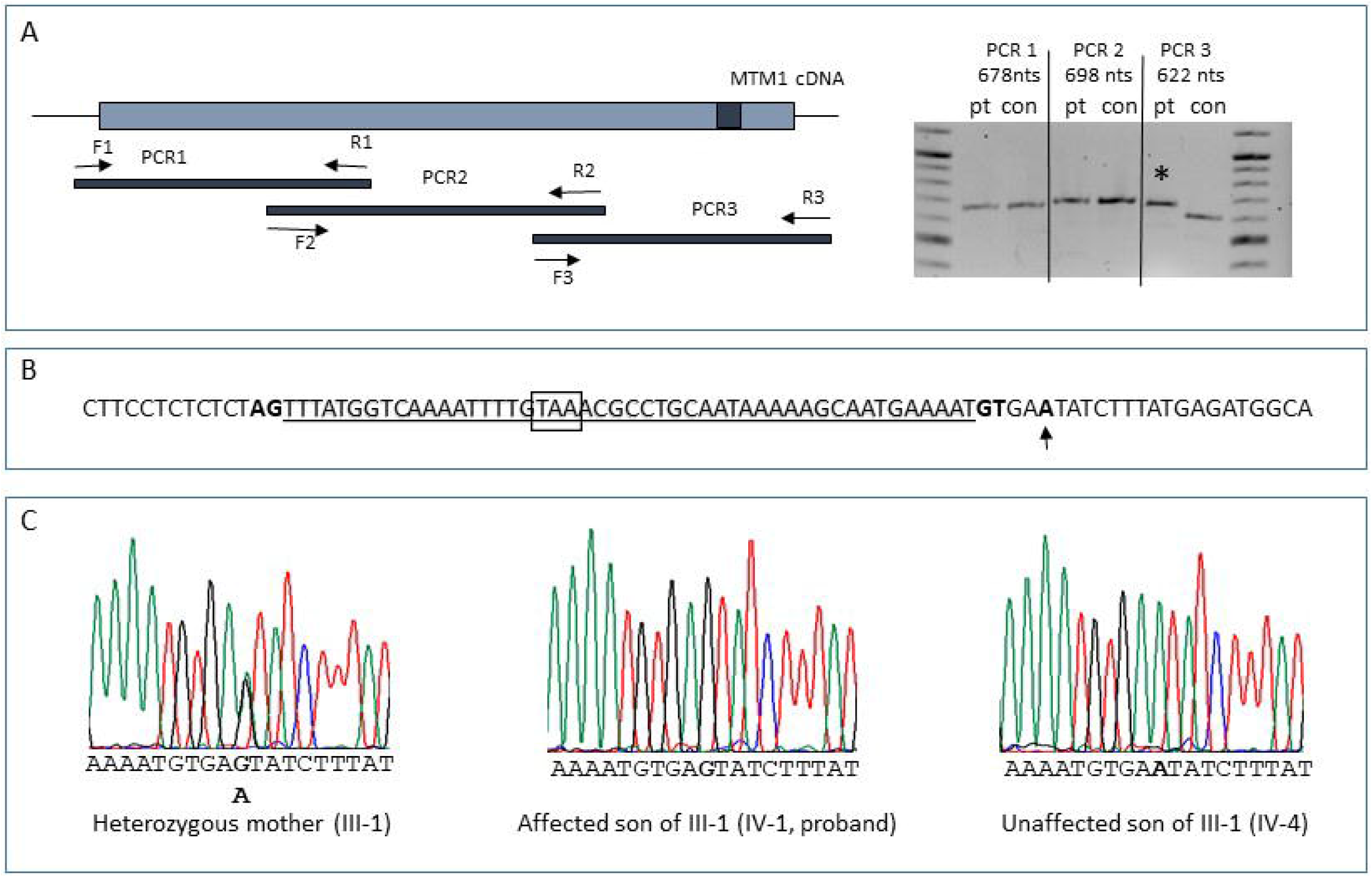
Identification of a pathogenic intron variant in *MTM1*. A, <u>Aberrant *MTM1* RNA splicing in the XLMTM1 family</u>. Three overlapping RT-PCR fragments encompassing the *MTM1* cDNA were designed based on published primers(Tosch et al. 2010). RNA from patient fibroblasts was isolated, reverse transcribed into cDNA and subjected to PCR using the three primer sets; F1/R1, F2/R2 and F3/R3. The right panel is a DNA agarose gel showing three RT-PCR products from patient (pt) and control (con) fibroblasts. PCR3 is larger in patient compared to control suggesting aberrant splicing and the inclusion of intronic sequence in the mature mRNA (pseudoexon). This is indicated by the black box in A. The expected size of PCR amplicons is shown above gel. Marker is 100bp ladder. B, <u>Genomic context of *MTM1* mutation</u>. The PCR3 band was sequenced and found to contain 48bp of *MTM1* intron 13 (underlined). Sequencing of genomic DNA in this intronic region showed the presence of a unique A>G substitution (in bold and indicated by arrow). The variant is 5 bp downstream of the 5’ cryptic splice site acceptor that is not normally activated. Canonical AG 3’ splice donor and GT 5’ splice acceptor sites that define the boundaries of the pseudoexon are in bold. The in-frame stop codon (TAA) is boxed. C, <u>Genotyping family members for the A>G variant in intron 13</u>. Family members were genotyped by sequencing across the genomic region containing the variant. A sequencing chromatogram shows that the carrier female (III-1) and mother to three of the affected males is heterozygous for the variant (left). Her affected son and proband (IV-1) is hemizygous for the G allele (middle) and an unaffected son (IV-4) is hemizygous for the A allele (right). DNA sequence of the variant is shown below each chromatogram and the variant alleles are in bold. Genotyping the remaining extended family members show that all affected males carry the G variant and all unaffected males the A variant.

### Variant results in altered splicing

*In silico* support for altered RNA splicing comes from Human Splicing Finder splice prediction software (http://www.umd.be/HSF3/HSF.shtml)(Desmet et al. 2009; Flicek et al. 2013). The G>A variant increases the MaxEnt splicing score from −0.4 to +7.8. With the threshold for splicing set at 3, the wild-type sequence is not predicted to be spliced and the mutant sequence is strongly predicted to be spliced (Fig. 3A). It is also noted that substitution of G for A at the +5 position creates a consensus sequence for a U1-snRNP splicing factor binding (A-G-G-U-A/G-A-**G**).

**Fig. 3:**
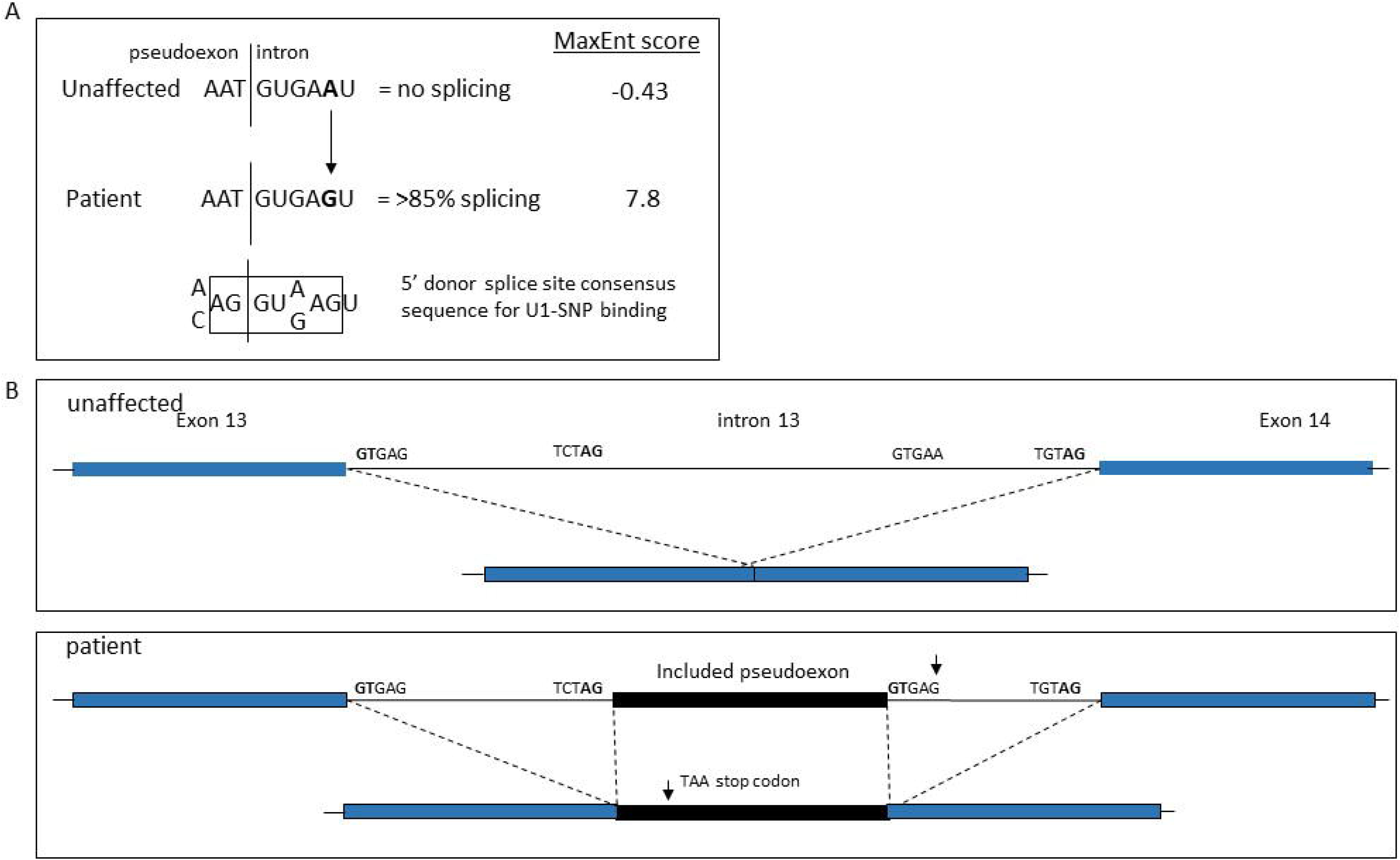
The A>G variant activates a cryptic 5’-splice site. A, <u>*In silico* analysis of altered splice site</u>. The A>G nucleotide change converts a pseudoexon, which is not normally spliced, into an ‘included pseudoexon’. Densitometric analysis of RT-PCR products indicates that 85% of *MTM1* transcripts contain the included pseduoexon. The 5’ss consensus sequence is shown below and the U1-snRNP binding site is boxed. Analysis using the MaxEnt tool of the Human Splicing Finder software predicts that the mutant sequence increases splicing efficiency. Since the score of the wild-type sequence is below the threshold of 3, it is not predicted to be spliced. In contrast, with a score of 7.8 the mutant sequence is strongly predicted to be spliced. B, <u>Schematic showing splicing pattern in patient and unaffected individuals due to A>G mutation</u>. Normal splicing pattern in unaffected individuals (top panel) with exclusion of intron 13 and splicing together of exons and 14. Intron splice site acceptor and donor sequences are shown. Splicing pattern of affected individuals (lower panel) showing the effect of A>G mutation (arrow). Activation of 5’ donor site (GTGAG) in pseudoexon also results in preferential use of existing 3’ acceptor site (TCTAG) at 5 end of pseudoexon to effect splicing of 48 bp pseudoexon sequence. Approximate location of TAA stop codon in included sequence is indicated.

Collectively, the experimental and prediction data indicate that the A>G variant activates a cryptic 5’-donor splice site in a pseudoexon in intron 13 leading to inclusion of intronic sequence in affected individuals. In unaffected individuals, the wild-type sequence is not sufficient to activate splicing of the pseudoexon and a normal splicing pattern is maintained. The new splice site acts in concert with the normal downstream 3’ acceptor splice site adjacent to exon 14. Similarly, the existing 5’ acceptor splice site near the start of the pseudoexon is used with the 5’ donor splice site adjacent to the upstream exon 13 (Fig 3B). The net result of the mutation is the preferential inclusion of the pseudoexon in individuals with the A/G variant.

### Decay of the mutant mRNA

The included 48bp pseudoexon sequence contains an in-frame TAA stop codon at chrX:149831295-149831297 suggesting that the mutant mRNA may undergo nonsense-mediated mRNA decay. This was confirmed by RT-PCR across the pseudoexon region in the proband and an affected brother that cells were available for (Fig. 4A). The pseudoexon band is much fainter in the affected sibling than in control fibroblasts. When patient fibroblasts were incubated with the translational inhibitor, cycloheximide, degradation of the mutant allele is halted, providing further evidence that the mutant allele undergoes nonsense-mediated mRNA decay. Quantitative PCR (qPCR) analysis reveals that the mutant allele is stabilized 7-fold in the presence of cycloheximide (Fig. 4B). Since the mutant allele is substantially degraded, we conclude that affected males produce no, or low levels, of the myotubularin-1 protein while carrier females produce sufficient myotubularin for normal development.

**Fig. 4:**
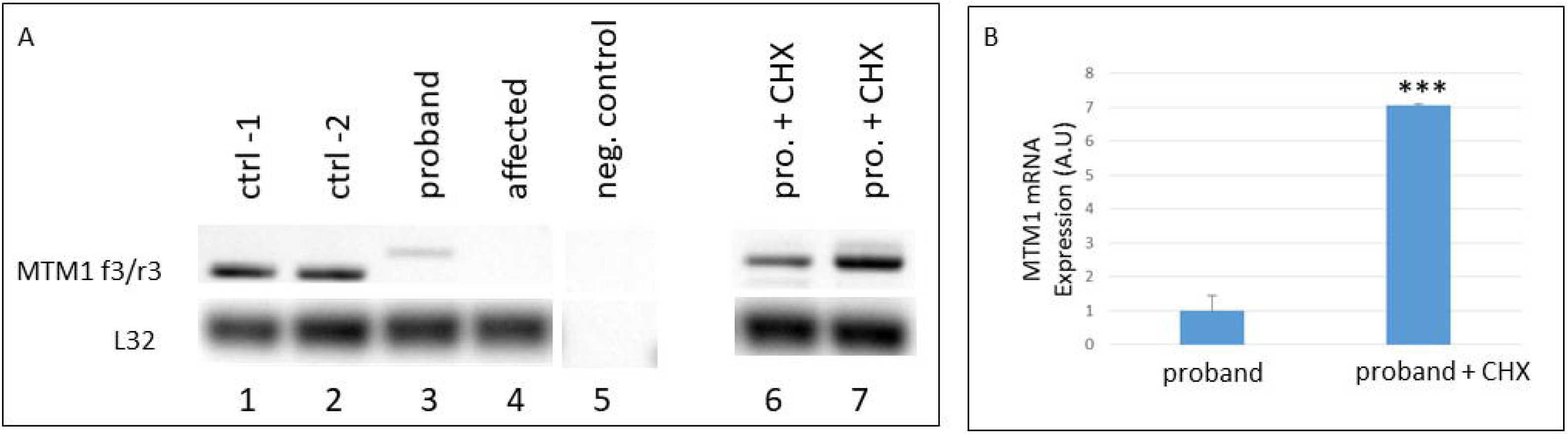
mRNA for mutant *MTM1* allele undergoes nonsense-mediated mRNA decay. A, <u>RT-PCR for normal and mutant mRNA</u>. cDNA samples from two control fibroblast cell lines (lanes 1 and 2), the proband (IV-1, lane 3) and an affected cousin (IV-6, lane 4) were amplified using the *MTM1* F3/R3 primers. No template negative control is shown in lane 5. Virtually no normally spliced *MTM1* mRNA is detected in patient cells compared to control cell, and low levels of mutant mRNA is present in patient cells indicating that the mutant allele is degraded. In a separate experiment, proband samples were cultured in the absence (lane 6) and presence (lanes 7) of cycloheximide. The mutant mRNA is stabilized by cycloheximide. L32 ribosomal mRNA is a housekeeping gene used as a control for cDNA. B, <u>qPCR analysis of cycloheximide-treated patient cells</u>. Quantitative PCR was conducted on cDNA isolated from proband cells treated with and without the translation inhibitor cycloheximide. Cycloheximide treatment results in 7x more mutant mRNA compared to untreated cells. ***P=0.0004, Students T-test.

## Discussion

We report a novel variant in intron 13 of the *MTM1* genes in a family with XLMTM. The mutation results in the inclusion of a pathogenic pseudoexon that leads to decay of the mutant mRNA. Affected males are essentially null for myotubularin-1, the *MTM1* gene product. The mutation is severe and three of the four affected males died in the neonatal period due to respiratory insufficiency while the fourth child survived only 14 months.

Tosch et al estimated that intronic mutations account for approximately 19% of *MTM1* mutations with the majority of these being intron-exon boundaries which are readily detectable by routine genomic DNA sequencing (Tosch et al. 2010). This group employed a screen specifically to detect intronic mutations in the *MTM1* gene and, in addition to identifying several novel intron-exon boundary mutations, identified the first pseudoexon mutation in *MTM1 (Tosch et al. 2010)*. RT-PCR and sequencing revealed 94 bp of included intron sequence in intron 7. The mutation created a functional splice acceptor site that when used together with an existing splice donor site resulted in exonization, leading to nonsense-mediated mRNA decay and loss of myotubilarin-1 protein. Our report is the second of these types of mutations in severe XLMTM1 and highlights the importance of deep intronic mutations in the pathogenesis of XLMTM1 and the need to include screening strategies when looking for *MTM1* mutations.

From a treatment perspective, the finding that carrier females develop normally yet express lower levels of myotubularin compared to females without *MTM1* mutations suggest that a modest increase in myotubularin levels may be therapeutically useful. It is unknown how much myotubularin-1 is required for normal development since carrier females produce 50% of the levels of normal males, assuming random X chromosome activation, yet appear phenotypically normal. This is relevant because one of the four affected males in the family who inherited the mutant allele survived past one year. Differences in cryptic splice site activation may explain this and there may be a threshold for myotubularin-1 levels such that the male that survived the neonatal period had less aberrant splicing and a higher incidence of normal splicing. One therapeutic approach may be to correct normal splicing in *MTM1* by the inhibition of pseudoexon activation in this family.

In summary, we described an intronic mutation which leads to 5’ splice site activation and the inappropriate inclusion of a segment of intron sequence into the mature mRNA. This leads to an in-frame premature stop codon and decay of the mutant mRNA which is catastrophic for the affected males in the family resulting in a lethal phenotype. Our study highlights the need to consider non-coding mutations in mutation screening in XLMTM1 families.

## Acknowledgements

We would like to thank the family who generously provided samples for this study.

## Statement of Ethics

This research was conducted ethically in accordance with the World Medical Association Declaration of Helsinki. Subjects provided their written informed consent and that the study protocol was approved by the Institutional Review Boards of Oregon Health and Science University (protocol #6758) and Henry Ford Hospital System (protocol #10967).

## Author Contributions

J.F. and D.B. were involved in the overall design of the project and conducted genetic analyses of samples. C.F. was the genetic counselor that liaised with the family and collected patient samples. P.D. conducted technical analyses. S.M. provided patient cell lines for the project. J.F., D.B., C.F. and S.M. wrote and edited the manuscript.

